# Initial approach to the Conservation Genetics of the Guatemalan Beaded Lizard (*Heloderma charlesbogerti*): A route to genetic rescue

**DOI:** 10.1101/2021.11.02.466675

**Authors:** Sergio González, Thomas Schrei, Brad Lock

## Abstract

In this study, samples from 33 Guatemalan Beaded Lizard (*Heloderma charlesbogerti*) were analyzed for genetic diversity. Twenty-three samples were obtained from wild individuals from two separate population areas, and 10 samples were obtained from captive individuals. Because the seasonally dry tropical forest habitat sampled for this study, is degraded and fragmented, it was hypothesized that beaded lizard populations were small and isolated and would be subject to genetic erosion and an elevated extinction risk. To test this hypothesis, eight microsatellite markers were employed to analyze 22 individual samples from the population of Cabañas, Zacapa, a single individual from the eastern-most population and 10 captive individuals of unknown origin. An average of three alleles per maker was reported for the Cabañas population, evidencing a low genetic diversity. In addition, a recent bottleneck event was detected and an effective population size of 19.6 was estimated. Demographic reconstruction using a Bayesian approach was inconclusive possibly due to a small dataset and shallow coalescence trees obtained with the generated data. No clear structuring pattern was detected for the Cabañas population and most samples from individuals in captivity were found to have similar alleles to the ones from Cabañas. Population designation is challenging without the genotyping of every wild population, but unique alleles were found in captive individuals of unknown origin that could suggest that different genotypes might exist within other, less studied, wild populations. Low genetic diversity, and a small effective population size represent a risk for the Cabañas population facing the threats of isolation, habitat loss and climate change. These findings suggest that genetic management of the Cabañas population might be utilized to avoid high rates of inbreeding and subsequent inbreeding depression.

## Introduction

This study presents current knowledge on the genetic diversity of the most well-known population of the endemic and endangered Guatemalan Beaded Lizard (*Heloderma charlesbogerti*). The loss of genetic diversity has been a central issue in the ongoing global crisis of biodiversity loss (Whiteley *et al*., 2014; Frankham *et al*., 2017). Human activities such as agriculture, cattle farming, increase in urban areas and roads, have led to wide-spread degradation and fragmentation of natural habitats, reducing the size of and connectivity between subdivided populations. These activities have also reduced migration rates and fragmented ideal habitat, which results in small and isolated populations that diverge and undergo bottleneck events that reduce genetic diversity (Tallmon *et al*., 2004; Frankham *et al*., 2017). Erosion of genetic diversity in these populations leads to higher inbreeding coefficients, stronger effects of genetic drift due to a lower effective population size (N_e_), and loss of adaptive potential which eventually can lead to population extirpation (Hughes *et al*., 2008; Frankham *et al*., 2010; Frankham *et al*., 2017). Therefore, Conservation Genetics has become an important tool for wildlife managers and scientists that are attempting to develop and implement effective conservation measures to ensure species survival (Tallmon, 2004; Höglund, 2009; Whiteley *et al*., 2014; Frankham *et al*., 2017).

Inbreeding depression is a phenomenon observed in populations where individuals with a high degree of relatedness, are breeding (Frankham *et al*., 2017). This leads to a loss of heterozygosity and a decrease in fitness usually attributed with the increase in genetic load of deleterious alleles and the expression of homozygous recessive alleles that are detrimental to the individual. Individuals with lower fitness are, therefore, less likely to survive to a reproductive age or successfully produce offspring. Effects at the population level also include a decreased population size and a decrease in overall fitness (Fredrickson *et al*. 2007; Ehlers *et al*., 2008; Funk *et al*., 2010; Johnson *et al*., 2010). The resulting positive feedback loop is referred to as an extinction vortex that can accelerate the decline of a population regardless of efforts to maintain the integrity of the habitat or increase the demographic population size (Höglund, 2009).

Genetic rescue has been proposed as a solution to mitigate or reverse these extinction vortices caused by inbreeding depression. This conservation action consists of enriching the genetic diversity of an endangered population by introducing individuals from other populations with a higher genetic diversity or different genotypes (Tallmon *et al*., 2004); which is expected to increase overall fitness. However, genetic rescues are rarely carried out in practice despite having produced promising results in cases such as that of the Florida panther, the European adder, and the Mexican wolf (Fredrickson *et al*., 2007; Ehlers *et al*., 2008; Johnson *et al*., 2010; Madsen *et al*., 2020). Most of the controversy regarding genetic rescue is centered on concerns of exogamic depression, or a decrease in fitness caused by the introduction of foreign individuals who might be adapted to different local conditions. However, this risk in practice, has been observed to be minimal. Even when some exogamic depression is initially observed (Whiteley *et al*.,2014), it is overcome in a few generations when fitness of the population recovers (Hwang *et al*., 2011).

The Guatemalan Beaded Lizard (*Heloderma charlesbogerti* Campbell and Vannini, 1988; Beck 2005; Reiserer et al. 2013; Fig. 1) is an endangered and endemic lizard species to Guatemala. It inhabits the Motagua Valley, in highly degraded and fragmented dry forest habitat, within four geographically delimited populations (Ariano-Sánchez and Salazar 2007; Anzueto and Campbell 2010). The total population for *H. charlesbogerti* has been estimated at 350 individuals (Ariano-Sánchez et al. 2014), although this figure is highly suspect due to limited region-wide sampling efforts. Population numbers of *H. charlesbogerti* have been reduced by habitat degradation, indiscriminate killing, and wildlife trafficking (Beck, 2005; Ariano-Sánchez *et al*., 2014).

**Figure 1.**
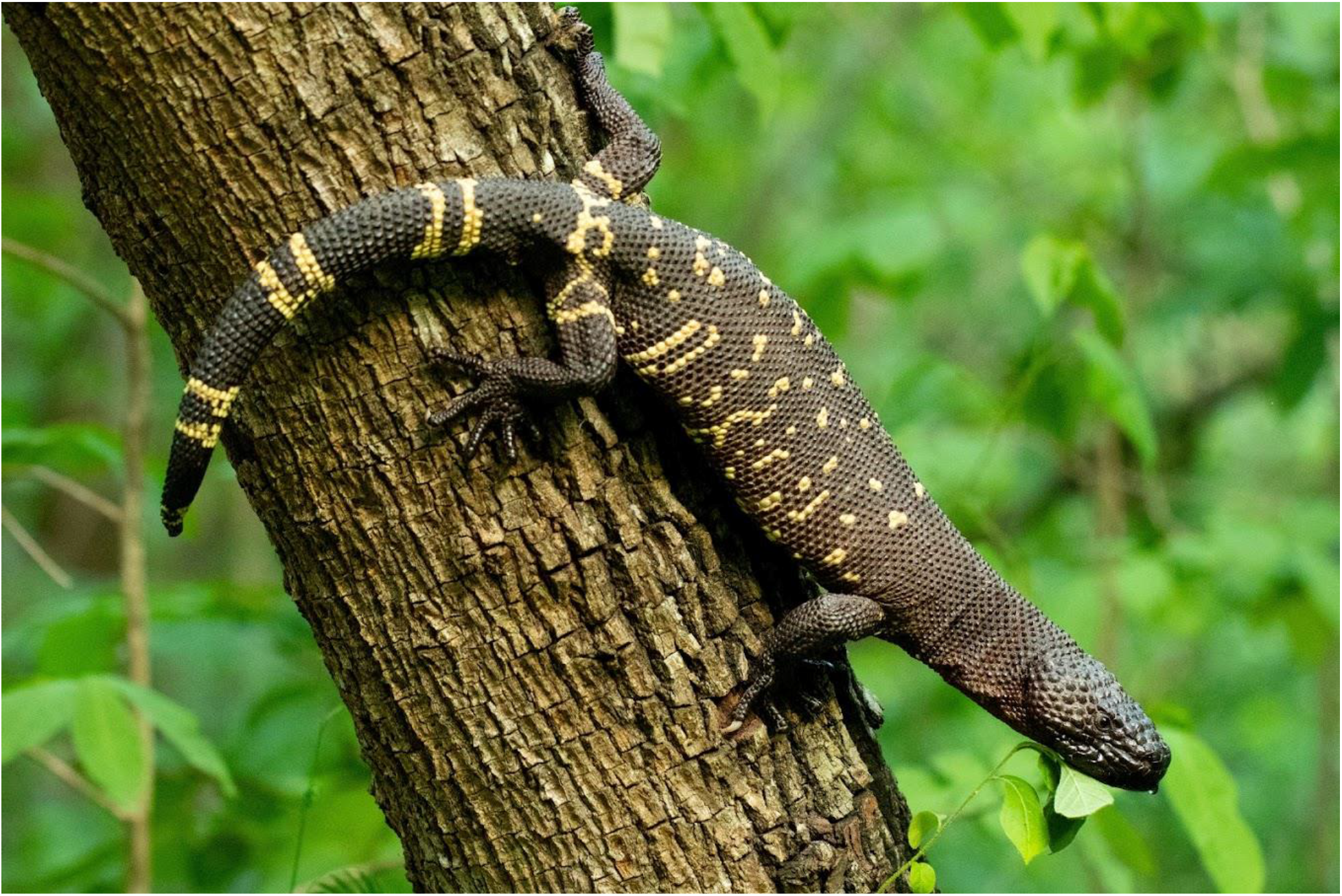
Individual of *Heloderma charlesbogerti* climbing a tree during its peak activity in the rainy season. Photograph: Thomas Schrei, 2021.

The habitat fragmentation and the small size of populations has raised suspicions of a possible bottleneck event on even the most preserved populations that could lead to inbreeding, a loss of genetic diversity and a subsequent depression of population fitness, making them susceptible to environmental fluctuations such as climate change and human pressure (Frankham et al. 2017; Ralls et al. 2018). A small effective population size, inbreeding depression and the impact of stochastic events that further degrade the genetic diversity of a small population may steer it into an extinction vortex (Höglund 2009). Climate change and the displacement of thermal niches in tropical lizards exacerbate the risks already presented (Huey et al. 2009; Sinervo et al. 2010), especially for *H. charlesbogerti*, as it exhibits a highly specific niche preference (Domínguez-Vega et al. 2012).

The importance of a genetic approach to the conservation of the species of *Heloderma* was explored by Douglas et al. (2010), pairing genetic diversity with phylogenetic history and geological processes affecting the evolution of helodermatid lizards. This helps explain the distribution and current taxonomic status and history of the species. Data from aa study on a finer scale must be employed to inform the development of more effective local action and management. This is the reason why the genetic diversity of each of the populations of *H. charlesbogerti* must be assessed. In this study, evaluation of the genetic diversity, genetic structure and demographic history of the Cabañas population and captive individuals using microsatellite markers, as used for vultures (Arshad et al. 2009), giant pandas (Lu et al. 2009), brown bears (Paetkau et al. 1998) and rattlesnakes (Bushar et al. 1998) is explored. Expected findings included a reduction in genetic diversity and a recent bottleneck event consistent with the habitat dynamics that the species has undergone in recent times due to human activities in regard to habitat distribution.

## Materials and Methods

### Study site

This study concentrated primarily on the dry forest habitat of the Motagua Valley, in mid-eastern Guatemala (Fig. 2). The main site of the study was the Cabañas population of *H. charlesbogerti*, found in Zacapa in the southwest department.

**Figure 2.**
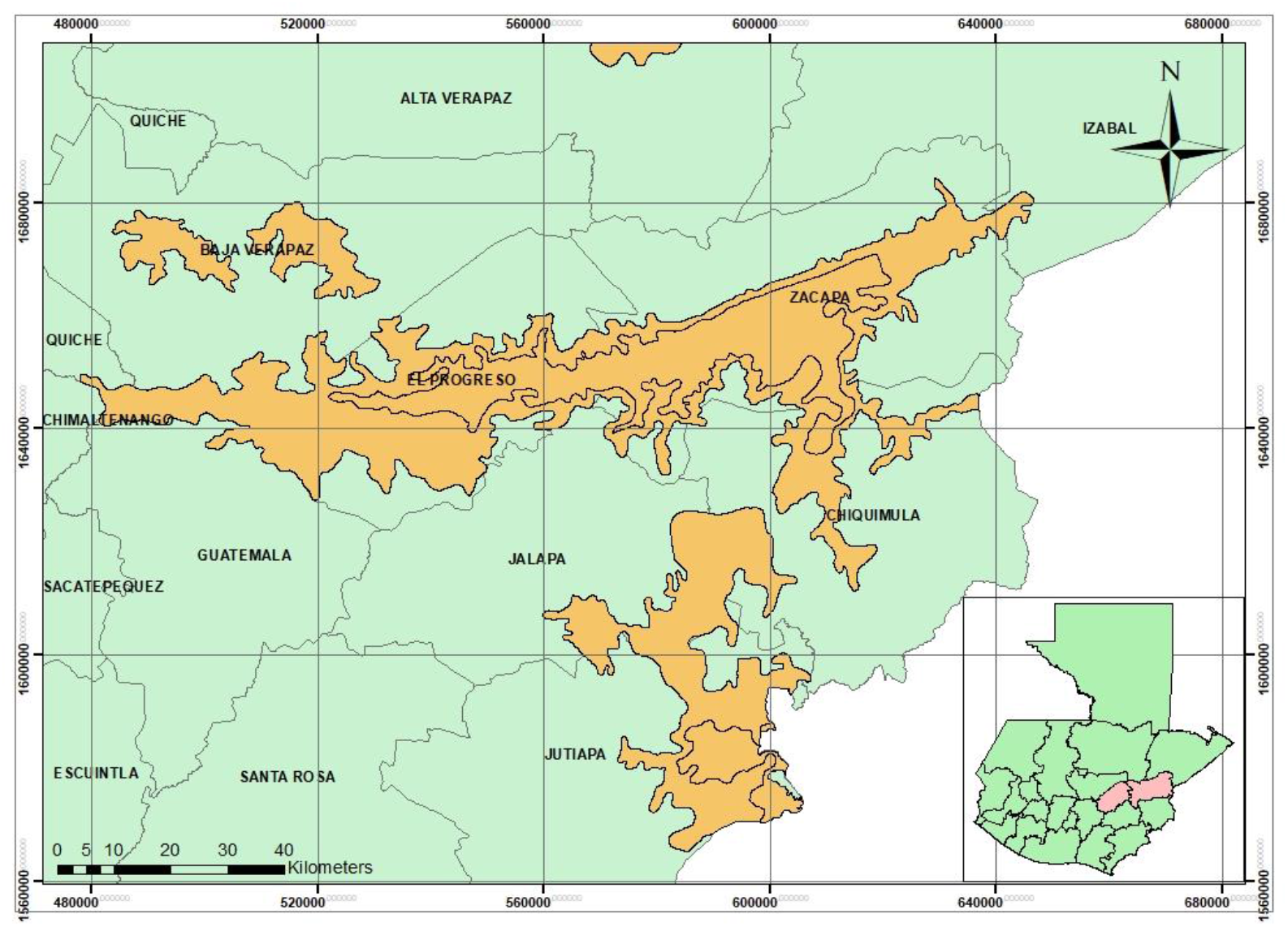
Arid regions of Eastern Guatemala, composed by spinous shrublands, dry subtropical and dry tropical forests. In the Motagua Valley these ecosystems are present in El Progreso and Zacapa (highlighted in the reference map). The grid shows UTM coordinates, datum WGS84 zone 16N.

### Sampling

A total of 33 samples (either moth swabs or blood samples) were collected and included in this study. Between 2014-2015, 22 individual lizards from the population of Cabañas, Zacapa, were sampled using mouth swabs (conserved in 95% ethanol until extraction). Between 2014-2015, 11 blood samples (1.0 ml each), preserved on FTA cards, were collected from the caudal tail vein, using a sterile, disposable syringe and a 22.0 gage needle from 10 captive lizards with unknown origin, and one wild individual from a separate, more eastern population.

### DNA Extraction

A modified phenol-chloroform DNA extraction using five 2-mm discs was used to obtain template DNA (González-Mollinedo 2018). The discs were put in a 1.5 mL tube with 500 μL of lysis solution (10 mM Tris-HCl, 100 mM NaCl, 10 mM EDTA and 2% SDS) and 20 μL of proteinase K (20 mg/mL). The tubes were incubated for four hours at 56ºC while shaking. One volume of phenol was added to each tube, thoroughly shaken, and centrifuged at 14,000 rpm for 10 minutes. The aqueous phase was retrieved and then washed with a volume of chloroform-isoamylic alcohol (24:1). The tubes were centrifuged at 14,000 rpm for 10 minutes and the aqueous phase was transferred to a tube holding 800 μL of isopropanol at −20ºC and 50 μL of ammonium acetate (3 M). The samples were stored overnight at −20ºC for precipitation. Afterwards, the tubes were centrifuged at 14,000 rpm for 30 minutes, at a temperature of 4ºC; the supernatant was discarded. The resulting pellet was washed with 800 μL of ethanol and centrifuged at 14,000 rpm for 20 minutes. The ethanol was discarded, and the tubes were laid open so that remnants of ethanol would evaporate. The pellet was suspended in 30 μL of warm TE solution.

The samples were amplified with 12 microsatellite markers developed by Hess et al. (2013) and 6 from Feltoon et al. (2005). The markers selected from Hess *et al*. (2013) were those which amplified with *H. horridum*, except for HESU018 which was discarded due to amplification problems. The PCR master-mix used was: Taq polymerase (Novagen NovaTaq 0.04U/μL), primers (0.5 μM each), amplification buffer (1X), MgCl2 (2.5 mM), dNTPs (0.2 mM for each nucleotide) DNA stock (50 ng).

### Sample Amplification and Allele Measurement

A Touchdown cycle was used to amplify the samples in a 100C Thermocylcer (Biorad), starting with an initial denaturalization at 95ºC, cycles of 95ºC denaturalization for 45 seconds, annealing temperatures from 60-51ºC, decreasing one degree every cycle, and an extension period of 45 seconds at 72ºC. After the decrease in annealing temperature reached 51ºC, another 30 cycles were run with a fixed annealing temperature at 51ºC. A final extension period of ten minutes at 72ºC finished the thermocycling process. At the end of the PCR, 10 μL of Stop Mix (0.2% w/v xylen-cyanol FF, 0.2% w/v bromophenol blue, 2% v/v EDTA 0.5 M, 3% distilled water and 95% v/v deionized formamide) were added to each product.

Amplification of the samples were confirmed using PAGE on 15 cm-long gels. The obtained products were then measured using denaturing urea PAGE on 50 cm-long gels as described by Summer *et al*. (2009). To measure the length of the amplified fragments, a Novagen 100-bp molecular weight ladder was loaded on three different lanes (on the sides and the middle of the gel) to compensate for the skewing of the sample lanes. The length of the fragments was calculated with the free software GelAnalyzer2010a.

### Data Analysis

Data analysis on the Cabañas population consisted of basic population genetic indicators using GenAlEx V. 6.503 (Peakall and Smouse 2006), discarding monomorphic markers. Number of alleles (*N_a_*),, Shannon Diversity Index (*I*), observed heterozygosity (*H_o_*), unbiased expected heterozygosity (*uH_e_*) and fixation coefficients (*F*) were calculated for the Cabañas population. Null alleles were checked for with FreeNA (Chapuis and Estoup, 2007). Effective population size (*N_e_*) was estimated using NeEstimator V. 2.01 (Do et al., 2014) with the Linkage Disequilibrium Method and with the minimum allele frequency of 0.01. Parametric and jackknife 95% confidence intervals are reported. Bottleneck V. 1.2.02 (Cornuet and Luikart 1997; Piry et al. 1999) was used to identify population bottleneck events on the Cabañas population using the Two-Phase Mutation Model (TPM) and the Stepwise Mutation Model (SMM). The sign test, the standardized differences test and the Wilcoxon sign test were considered. TPM parameters for proportion of SMM in TPM= 0.0000 and for variance in geometric distribution for TPM= 0.36. The simulation ran for 1,000 iterations.

An approximate Bayesian computation approach was used to reconstruct the demographic history of the Cabañas population using DIYABC V. 2.1.0 (Cornuet et al., 2014). The population was assumed to be isolated during the reconstruction and three scenarios were simulated: 1) Constant population size, 2) a bottleneck event and 3) an ancient contraction in population size consistent to the fragmentation of seasonally dry forests during the Quaternary glacial periods (Pennington et al., 2000), a recovery, and a recent bottleneck event. The three scenarios were simulated with increasingly constraining priors. The best set of priors were chosen due to their likeness to the proposed scenarios in reality, and the clarity of the posterior distributions generated. Default mutation rates for microsatellites were used, since there is no information for the mutation rates of the markers used in the studied species (see Table S1 for full set of priors and scenarios). Each scenario was simulated 1,000,000 times to generate posterior distributions. The probability of each scenario was calculated with the posterior distributions of the simulated datasets and the confidence of the scenario comparison was calculated with 1,000 pseudo-observed data sets where the best scenario was obtained.

Further analysis considered individuals in captivity. Genetic structure was inferred considering the whole dataset using the software STRUCTURE V. 2.3.4 (Pritchard *et al*., 2000). Simulations were run with the Admixture model for possible populations (*K*) values of 1-3, with a burn-in period of 100,000 iterations and 1,000,000 MCMC interactions. The most probable value of *K* was obtained using Structure Harvester with an input of four runs for each value of *K* (Earl and von Holdt 2011).

## Results

### Population genetics for Cabañas

Results for the Cabañas population are shown in Table 1. Markers with monomorphic results and those with null alleles were eliminated from the analysis. The mean alleles per marker were 3 (Standard Error: 0.378, Table 1), where marker HESU004 was the most polymorphic with 5 alleles and the highest diversity index. The population seems to have a low genetic diversity across polymorphic markers. The population also shows *H_o_*<*H_e_* and strongly negative fixation coefficients (F_IS_) for all markers, with a mean of −0.255. Effective populations size was estimated to be 19.6 individuals (95% CI: 6.9-433.1. When using the jackknife for the samples, the 95% CI is: 6.3-∞).

**Table 1.** Allele number (N_a_), effective allele number (N_e_), diversity index (I), observed (H_o_) and unbiased expected heterozygosity (uH_e_) and fixation coefficients (F) for the Cabañas population across analyzed SSR markers. Markers with null alleles were removed. Negative F values denote an excess of heterozygosity in comparison to the expected.

| Locus | N | $N_a$ | $I$ | $H_o$ | $uH_e$ | $F$ |
| --- | --- | --- | --- | --- | --- | --- |
| HESU004 | 20 | 5.000 | 1.395 | 0.850 | 0.740 | -0.179 |
| HESU007 | 18 | 3.000 | 0.868 | 0.722 | 0.565 | -0.315 |
| HESU008 | 22 | 3.000 | 0.906 | 0.864 | 0.566 | -0.563 |
| HESU009 | 20 | 3.000 | 0.819 | 0.600 | 0.477 | -0.290 |
| HESU010 | 22 | 4.000 | 1.226 | 0.955 | 0.695 | -0.406 |
| HELO1 | 22 | 2.000 | 0.562 | 0.500 | 0.384 | -0.333 |
| HELO2 | 22 | 2.000 | 0.625 | 0.455 | 0.444 | -0.048 |
| HELO3 | 22 | 2.000 | 0.642 | 0.409 | 0.460 | 0.090 |
| Mean | 21.000 | 3.000 | 0.880 | 0.669 | 0.541 | -0.255 |
| SE | 0.535 | 0.378 | 0.105 | 0.073 | 0.044 | 0.073 |

Bottleneck simulations were run for the Cabañas population and a recent bottleneck event was indicated. Statistically significant p-values were found for heterozygosity excess in all tests except for the Sign test for SMM. The simulation results are summarized on Table 2. The Bayesian demographic inference did not yield a clear best scenario and it could not distinguish between the scenarios with population contractions and the one with constant effective population size as confidence intervals overlapped in all sets of priors evaluated and the selected set of priors showed a confidence of 0.504 in the scenario selection favoring the one with constant population size. The DIYABC analysis can be found in Supplementary Material.

**Table 2.** Summary of *p*-values produced by Bottleneck simulation for the Cabañas population. Statistically significant tests for bottlenecks were found when testing for heterozygote excess (significant p-values are highlighted in bold).

| Mutation Model | Wilcoxon Test |  | Sign test |  | Mode Shift |
| --- | --- | --- | --- | --- | --- |
|  | One-tailed Hx | One-tailed Hd | Hd:Hx | p-value |  |
| TPM | **0.002\*** | 1.000 |  | **0.005\*** | No |
| SMM | **0.004\*** | 0.998 |  | 0.071 |  |

### Genetic structure

Structure Harvester determined that the most adequate number of clusters was two (*K*=2). The individuals analyzed were given a Q value and were arranged accordingly (Fig. 3). The analysis did not show clear signs of structural differentiation amongst the individuals from Cabañas, the individual from the other wild population or the ones in captivity. When the individuals in captivity and the single wild individual from the eastern population were assigned to different populations, the mean F_ST_ value reported was 0.233, but the population of origin for most of the individuals in captivity cannot be confirmed.

**Figure 3.**
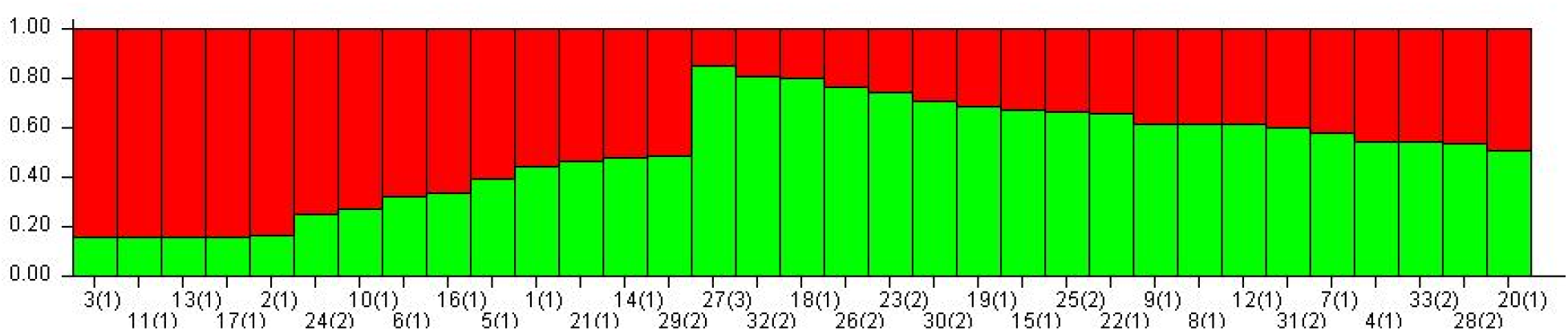
Population structure analysis performed by Structure V. 2.3.4 arranging individuals by its Q scores. Population assignment is represented inside parenthesis, Pop. 1= Cabañas, Pop. 2= Captivity, Pop. 3= other population. Structure Harvester established *K*=2 and each subgroup are represented by a different color. Note that even though the assignment for some individuals is clearly to one or the other group, they come from the same population.

### Unique alleles

Individuals in captivity could not be analyzed as a stand-alone population and the data that could be generated was not useful because they could not be assigned to any known population. However, unique alleles were found in at least four individuals in captivity and in the individual from the easternmost population that did not match the predominant alleles in Cabañas for markers HESU004 and HESU007.

## Discussion

*H. charlesbogerti* is an endemic species with a limited distribution with the majority of known habitat being severely fragmented and degraded. The seasonally dry tropical forest in Guatemala has no formal conservation status and is commonly perceived neglected as dry shrubland that requires little attention rather than as an important forest type (Janzen, 1988). This lack of government protection along with unregulated human intervention and cultural dislike for the species, has reduced gene flow along the distribution of the species.

Methodological limitations of this study include the lack of access to technology like capillary electrophoresis to be able to estimate microsatellite sizes which might be more replicable than the urea-PAGE we used or microsatellite markers that were designed for a different species. Sample size might seem low, but we managed to obtain tissues from approximately one fourth of the estimated population size for Cabañas and provided initial data for both the conservation genetics of the species and preliminary information to ongoing conservation efforts. Attempts to reconstruct the demographic history of the Cabañas population were inconclusive with the current markers, as Bayesian approaches benefit from a larger sample size and a larger number of highly polymorphic markers to achieve robust coalescence; In that regard, the data generated is lacking. Low allele diversity found in the markers and the differences of one or two repeats between alleles in this study also suggest a shallow coalescence tree which could not be useful for Bayesian analysis of demographic processes. Nonetheless, the bottleneck event found using the heterozygote excess tests seems likely, since it is sensitive to recent bottleneck events and the population states of excess heterozygosity are transient after significant reductions in effective population size (Cornuet & Luikart, 1996; Luikart et al., 1998).

The status of the population can be assessed in comparison to similar populations with available information about their genetic diversity. Farrar *et al*. (2017) carried out a similar study for a population of *H. suspectum* with a similar population size calculated by mark-recapture methods to the population of *H. charlesbogerti* in Cabañas using the same microsatellite markers in this study. They reported a population size of 80 individuals using mark-recapture methods and an effective population size of 94 individuals (95% confidence interval: 80.7-111.2). In this study, an effective population size for Cabañas of 19.6 individuals (95% CI: 6.9-433.1) was determined. Previous monitoring of this population shows that there are a minimum of 80 lizards that have been individually identified with transponder chips for Cabañas and a recent parallel mark-recapture study estimated a population size of approximately 160 (Schrei, unpublished data). Even with a larger estimated population size, genetic diversity is much lower in Cabañas, with a mean of 3 alleles per marker in comparison to the 8.06 that Farrar *et al*. (2017) reported.

A low effective population size accelerates inbreeding in small and isolated populations and an inbreeding coefficient of 0.25 could be reached in 11 generations, when the population might start presenting heterozygosity deficiency when considering an F_IS_ value of −0.255 and assuming that heterozygosity would decrease proportionally (generational inbreeding rate; Frankham *et al*., 2017). The excess heterozygosity could be evidence of a recent contraction of the population, but it can also be an artifact of a small population size or a consequence of differences of allele frequencies between males and females in a small population (Pudovkin et al., 1996). No large differences in allele frequencies between sexes were found in our dataset, but we cannot be confident in this result since there are very few alleles per marker to be able to be certain of the distribution of genetic diversity among sexes. An alternative explanation for this paradoxical heterozygosity excess and low genetic diversity could be that the population has undergone purging of highly deleterious genotypes that might have been predominantly homozygous (Byers and Waller 1999; Crnokrak and Barrett 2002; García-Dorado 2012; López-Cortegano et al. 2018). Despite the purging of deleterious genetic load has been mostly discussed for plants populations that are capable of self-fertilization and expose homozygous genotypes to selective forces (Byers and Waller, 1999), examples in severely bottlenecked animal populations are plausible (Grossen et al. 2020). Low population sizes and inbreeding expose the deleterious genetic load to selection and the individuals with such genotypes would probably not be viable in the population at variable stages of life, from the viability of the eggs produced to the probability of reproductive success of adult individuals (García-Dorado, 2012; Frankham et al., 2017). However, more information on the fitness of individuals in the population, and information from genomic markers and different analyses would be needed to confirm that this is the case. This hypothesis remains as conjecture with the current data. Nonetheless, our results indicate that given the low population estimate and effective population size, the Cabañas population of beaded lizards is at risk of being driven into inbreeding depression and an extinction vortex, even under the protection of a conservation program (Höglund 2009). This would make the population especially vulnerable to climate change, desertification, and other catastrophic scenarios (Domínguez-Vega et al. 2012; Huey et al. 2009; Sinervo et al. 2010).

In this study, only eight polymorphic markers were identified. One reason for this limited result may be that this study used primers designed for *H. suspectum* (Feltoon et al. 2007; Hess et al. 2013). However, despite this limitation we believe that the information generated is relevant. We have identified a reduced genetic diversity and reduced effective population size likely secondary to a very recent population bottleneck. This finding alone, is enough to consider the need for genetic management of this species (Frankham *et al*., 2017). To better inform this proposed genetic rescue program, genotyping of the other populations that might as well have experienced bottleneck events and loss of genetic diversity is necessary. This possibility is supported by the identification of unique alleles from individuals in captivity indicating that other populations might retain gene pools that could help increase genetic diversity and avoid inbreeding if gene flow is restored into the Cabañas population. The alleles shared across all individuals suggests that gene flow occurred at some point and that it might have been interrupted recently by human activities that might have also contributed to the bottleneck event in Cabañas. Finally, genotyping other populations of the species might help to assign a population of origin for the individuals in captivity which currently are of uncertain or unknown origin, but due to their genotypes, we believe that most of them originated from the Cabañas population. Six of the animals held in captivity were collected by foreign researchers when they worked in the country. One of those animals is said to come from La Unión, Zacapa, which appears to be the most isolated location where the species has been reported and we believe that the most genetically distinct of the individuals that were collected by them may have come from this site.

Further research and sampling efforts must go into genotyping the remaining populations of the species and this information will aid in designing a genetic rescue strategy that could bring adaptative potential and resilience to the species (Ralls et al. 2018). The need for new markers is also pressing since current microsatellite markers have not been able to generate robust datasets. Genomic approaches such as RADseq or genome sequencing can help increase the number of informative markers required to reconstruct the demographic history of the populations of the species and test scenarios of population isolation and contraction. Breeding programs for the species would benefit from the genotyping of the individuals that are being bred and estimations of kinship would ensure that the best mating pairs are selected to maintain genetic diversity. Establishing breeding programs without the pertinent genetic information may be counterproductive. Furthermore, studies of the effects of low genetic diversity, low effective population size and isolation of the populations of the species should be carried out and possibly revisit certain aspects of its natural history that might have been taken for granted. Interpopulation crosses could reveal higher performance in key traits that reflect fitness, and this could also contribute to the conservation of the species. The preliminary results generated in this study have already been incorporated in the National Conservation Strategy for the species and the urgency of genetic rescue should be strongly considered in conservation efforts in the coming years.

## Supporting information

Microsatellite data

## Acknowledgments

We would like to thank Zootropic, La Aurora Zoo, Antigua Exotic and the National Museum of Natural History in Guatemala for their help in obtaining useful samples for this project. We thank the Biology Department of Universidad del Valle de Guatemala for their support. We thank Julia Lesher, Anotnio Solé, Per Pallsbøl and Renaud Vitalis for their valuable help with data analysis. We thank the Georgia Institute of Technology and Zoo Atlanta for their financial support to carry out this project.

## Supplementary Material

Bayesian computation was used to try to reconstruct the demographic history of the Cabañas populations. Three different scenarios and three different sets of priors were used to attempt this reconstruction, but neither set of priors allowed to identify the most likely scenario for the population (Table S1). We argue that the lack of diversity in the markers used produces a shallow coalescence tree that makes the inference of population contractions difficult. This method contrasts with the Bottleneck tests that rely on the observed and expected heterozygosity in the dataset to infer the probability of population bottlenecks. DIYABC could perform better with a larger number of polymorphic markers to infer the demographic history of the Cabañas population.

**Table S1.** Prior sets used to infer the demographic history of the Cabañas population, the prior assumptions for each set and the confidence in the choice of scenario generated by each set. None of the sets of priors allowed to clearly infer a demographic history for the population.

| Set 1 |  |  |  |  |
| --- | --- | --- | --- | --- |
| Scenario | Parameters | Value | Prior assumptions | Confidence of scenario choice |
| Constant population size | N1 | 10.0-4000.0 | NA | 0.504 |
| Recent bottleneck | N1 | 10.0-4000.0 | Na>N1 |  |
|  | T2 | 10.0-5000.0 |  |  |
|  | Na | 1000.0-10000.0 |  |  |
| Ancient and recent bottleneck | N1 | 10.0-4000.0 | Na>N1, Nb<Na, t3>t2, Nc>Nb |  |
|  | T2 | 10.0-5000.0 |  |  |
|  | Na | 1000.0-10000.0 |  |  |
|  | T3 | 3000.0-10000.0 |  |  |
|  | Nb | 100.0-10000.0 |  |  |
|  | t4 | 10000.0-20000.0 |  |  |
|  | Nc | 1000.0-50000.0 |  |  |
| Set 2 |  |  |  |  |
| Scenario | Parameters | Value | Prior assumptions | Confidence of scenario choice |
| Constant population size | N1 | 1.0-4000.0 | NA | 0.503 |
| Recent bottleneck | N1 | 10.0-4000.0 | Na>N1 |  |
|  | T2 | 10.0-5000.0 |  |  |
|  | Na | 1.0-10000.0 |  |  |
| Ancient and recent bottleneck | N1 | 10.0-4000.0 | Na>N1, Nb<Na, t3>t2, Nc>Nb |  |
|  | T2 | 10.0-5000.0 |  |  |
|  | Na | 1000.0-10000.0 |  |  |
|  | T3 | 3000.0-10000.0 |  |  |
|  | Nb | 10.0-10000.0 |  |  |
|  | t4 | 10000.0-20000.0 |  |  |
|  | Nc | 1000.0-50000.0 |  |  |
| Set 3 |  |  |  |  |
| Scenario | Parameters | Value | Prior assumptions | Confidence of scenario choice |
| Constant population size | N1 | 10.0-4000.0 | NA | 0.52 |
| Recent bottleneck | N1 | 10.0-4000.0 | NA |  |
|  | T2 | 10.0-5000.0 |  |  |
|  | Na | 1000.0-10000.0 |  |  |
| Ancient and recent bottleneck | N1 | 10.0-4000.0 | Nb<Na, Nc>Nb |  |
|  | T2 | 10.0-5000.0 |  |  |
|  | Na | 1000.0-10000.0 |  |  |
|  | T3 | 3000.0-10000.0 |  |  |
|  | Nb | 100.0-10000.0 |  |  |
|  | t4 | 10000.0-20000.0 |  |  |
|  | Nc | 1000.0-50000.0 |  |  |

## Notes

### Competing Interest Statement

The authors have declared no competing interest.

## References

Anzueto, R, Campbell JA (2010) Guatemala Beaded Lizard (*Heloderma charlesbogerti*) on the Pacific Versant of Guatemala. Southwest Nat 55: 453–454.

Ariano-Sánchez D, Salazar G (2007) Notes on the Distribution of the Endangered Lizard, *Heloderma charlesbogerti*, in the Dry Forests of Eastern Guatemala: An Application of Multi-criteria Evaluation to Conservation. Iguana 14: 152–158.

Ariano-Sánchez, D, Muñoz-Alonso A, Márquez LC, Acevedo M (2014) Heloderma horridum. The IUCN Red List of Threatened Species 2014. http://dx.doi.org/10.2305/IUCN.UK.2014-1.RLTS.T9864A3152367.en. Accesed 31 January 2018.

Arshad M, Pedall I, González J et al (2009) Genetic Variation of Four Gyps Species (*Gyps bengalensis, G. africanus, G. indicus* and *G. fulvus*) based on Microsatellite Analysis. J Raptor Res 43: 227–236

Beck DD (2005) Biology of Gila Monsters and Beaded Lizards. University of California Press, Berkley and Los Angeles.

Bushar LM, Reinert HK, Gelbert L (1998) Genetic Variation and Gene Flow within and between Local Populations of the Timber Rattlesnake, *Crotalus horridus*. Copeia 1998: 411–422.

Byers, D. L. and D. M. Waller. 1999. Do Plant Populations Purge Their Genetic Load? Effects of Population Size and Mating History on Inbreeding Depression. Annual Review of Ecology and Systematics 1999 30(1): 479–513

Campbell JA, Vannini JP (1988) A New Subspecies of Beaded Lizard, *Heloderma horridum*, from the Motagua Valley of Guatemala. J Herpetol 22: 457–468.

Chapuis, M.P., and A. Estoup. 2007. Microsatellite null alleles and estimation of population differentiation. Mol. Biol. Evol. 24(3): 621–631.

Cornuet JM, Luikart G (1996) Description and power analysis of two tests for detecting recent population bottlenecks from allele frequency data. Genetics 144: 2001–2014.

Cornuet JM, Pudlo P, Veyssier J, Dehne-Garcia A, Gautier M, Leblois R, Marin JM, Estoup A. 2014. DIYABC v2.0: a software to make approximate Bayesian computation inferences about population history using single nucleotide polymorphism, DNA sequence and microsatellite data. Bioinformatics, 30(8): 1187–1189, https://doi.org/10.1093/bioinformatics/btt763

Crnokrak, P. and Barret, S. C. H. 2002. Perspective: Purging the Genetic Load: A Review of the Experimental Evidence. Evolution, 56(12): 2347–2358.

Do, C., Waples, R. S., Peel, D., Macbeth, G. M., Tillett, B. J. & Ovenden, J. R. (2014). NeEstimator V2: re-implementation of software for the estimation of contemporary effective population size (Ne) from genetic data. Molecular Ecology Resources. 14, 209–214.

Domínguez-Vega H, Monroy-Vilchis O, Balderas-Valdivia CJ et al (2012) Predicting the potential distribution of the beaded lizard and identification of priority areas for conservation. J Nat Conserv 20: 247–253.

Douglas ME, Douglas MR, Schuett GW et al (2010) Conservation phylogenetics of helodermatid lizards using multiple molecular markers and a supertree approach. Mol Phylogenet Evol 55:153–167.

Earl DA, vonHoldt BM (2011) STRUCTURE HARVESTER: a website and program for visualizing STRUCTURE output and implementing the Evanno method. Conservation Genet Resour 4: 359–361.

Ehlers A, Worm B, Reusch TB (2008) Importance of genetic diversity in eelgrass *Zostera marina* for its resiliance to global warming. Mar Ecol Prog Ser 355: 1–7.

Farrar VS, Edwards T, Bonine KE (2017) Elusive does not always equal rare: genetic assessment of a protected Gila monster (*Heloderma suspectum*) population in Saguaro National Park, Arizona. Amphibia-Reptilia 38: 1–14.

Feltoon AR, Lehn C, Guerra T et al. (2007) Isolation and characterization of six microsatellites in the Mexican beaded lizard *Heloderma horridum*. Mol Evol Notes 7: 433–435.

Frankham R, Ballou JD, Briscoe DA (2010) Introduction to Conservation Genetics. 2^nd^ edn. Cambridge University Press, Cambridge.

Frankham R, Ballou JD, Ralls K et al. (2017) Genetic Management of Fragmented Animal and Plant Populations. Oxford University Press, Oxford.

Fredrickson RJ, Siminski P, Woolf M et al. (2007) Genetic rescue and inbreeding depression in Mexican wolves. Proc Biol Sci 274: 2365–2371.huey

Funk WC, Forsman ED, Johnson M et al. (2010) Evidence of recent population bottlenecks in Northern Spotted Owls (*Strix occidentalis caurina*). Conserv Genet 11: 1013–1021.

García-Dorado, A. 2012. Understanding and Predicting the Fitness Decline of Shrunk Populations: Inbreeding, Purging, Mutation, and Standard Selection. Genetics, 190(4): 1461–1476.

González-Mollinedo S. 2018. Protocol for DNA extraction from Helodermatid blood samples preserved on FTA cards. Mesoamerican Herpetology 5(1): 197–198.

Grossen, C., Guillaume, F., Keller, L.F. et al. 2020. Purging of highly deleterious mutations through severe bottlenecks in Alpine ibex. Nat Commun 11, 1001. https://doi.org/10.1038/s41467-020-14803-1

Hess MR, Edwards T, Edmunds DA et al (2013) Characterization of STR/microsatellite primers for the Gila monster, *Heloderma suspectum* screened from paired-end Illumina shotgun sequencing. Conserv Genet Resour 5:1121–1123.

Höglund J (2009) Evolutionary Conservation Genetics. Oxford Scholarship Online, Oxford.

Huey RB, Deutsch CA, Tewksbury JJ et al. (2009) Why tropical forest lizards are vulnerable to climate warming. Proc Biol Sci 276: 1939–1948.

Hughes AR, Inouye BD, Johnson MT et al (2008) Ecological consequences of genetic diversity. Ecol Lett 11: 609–623.

Hwang, A.S., Northrup, S.L., Alexander, J.K. et al. Long-term experimental hybrid swarms between moderately incompatible Tigriopus californicus populations: hybrid inferiority in early generations yields to hybrid superiority in later generations. Conserv Genet 12, 895–909 (2011). https://doi.org/10.1007/s10592-011-0193-1

Johnson WE, Onorato DP, Roelke ME et al. (2010) Genetic restoration of the Florida panther. Science 329: 1641–1645.

López-Cortegano, E., Bersabé, D., Wang, J. and García-Dorado, A. 2018. Detection of genetic purging and predictive value of purging parameters estimated in pedigreed populations. Heredity, 121(1): 38–51.

Luikart G, Allendorf FW, Cornuet JM et al (1998) Distortion of allele frequency distributions provides a test for recent population bottlenecks. J Heredity 89: 238–247.

Madsen, T, Loman, J, Anderberg, L, Anderberg, H, Georges, A, Ujvari, B (2020) Genetic rescue restores long-term viability of an isolated population of adders (*Vipera berus*). Current Bioloy, 30(21): R1297–R1299

Paetkau D, Waits LP, Clarkson PL et al (1998) Variation in Genetic Diversity across the Range of North American Brown Bears. Conserv Biol 12: 418–429.

Peakall R, Smouse PE (2006) GenAlEx 6: genetic analysis in Excel. Population genetic software for teaching and research. Mol Ecol Notes 6: 288–295.

Piry S, Luikart G, Cornuet JM (1999) BOTTLENECK: A Computer Program for Detecting Recent Reductions in the Effective Population Size Using Allele Frequency Data. J Heredity 90: 502–503.

Pritchard JK, Stephens M, Donelly P (2000) Inference of population structure using multilocus genotype data. Genetics 155: 945–959.

Pennington, R. T., Prado, D. & Pendry, C. (2000). Neotropical seasonally dry forests and Quaternary vegetation changes. Journal of Biogeography 27: 261–273.

Pudovkin, A I, D V Zaykin, D Hedgecock, On the Potential for Estimating the Effective Number of Breeders From Heterozygote-Excess in Progeny, Genetics, Volume 144, Issue 1, 1 September 1996, Pages 383–387, https://doi.org/10.1093/genetics/144.1.383

Ralls K, Ballou JD (2004) Genetic status and management of California condors. The Condor 106:215–228.

Ralls, K., Ballou, J.D., Dudash, M.R., Eldridge, M.D.B., Fenster, C.B., Lacy, R.C., Sunnucks, P. and Frankham, R. (2018), Call for a Paradigm Shift in the Genetic Management of Fragmented Populations. CONSERVATION LETTERS, 11: e12412. https://doi.org/10.1111/conl.12412

Reiserer R, Schuett GW, Beck DD (2013) Taxonomic reassessment and conservation status of the beaded lizard, *Heloderma horridum* (Squamata: Helodermatidae). Amphib Reptile Conserv 7: 74–96.

Sinervo B, Méndez-de-la-Cruz F, Miles DB et al. (2010) Erosion of Lizard Diversity by Climate Change and Altered Thermal Niches. Science. 328: 894–899.

Summer H, Grämer R, Dröge P (2009) Denaturing Urea Polyacrylamide Gel Electrophoresis (Urea PAGE). J Vis Exp 2009: 1485.

Tallmon DA, Luikart G, Waples RS. (2004) The alluring simplicity and complex reality of genetic rescue. Trends Ecol Evol 19: 489–496.

Whiteley, AR, Fitzpatrick, SW, Funk, WC, Tallmon, DA (2014) Genetic rescue to the rescue. Trends in Ecology and Evolution, 30(1): 42–49.

